# Automated Cell Type Annotation with Reference Cluster Mapping

**DOI:** 10.1101/2024.11.30.626130

**Authors:** Valerio Galanti, Lingting Shi, Elham Azizi, Yining Liu, Andrew J. Blumberg

## Abstract

Single-cell RNA sequencing has transformed the field of cellular biology by providing unprecedented insights into cellular heterogeneity. However, characterizing scRNA-seq datasets remains a significant challenge. We introduce RefCM, a novel computational method that combines optimal transport and integer programming to enhance the annotation of scRNA clusters using established reference datasets. Our method produces highly accurate cross-technology, cross-tissue, and cross-species mappings while remaining tractable at atlas scale, outperforming existing methods across all these tasks. By providing precise annotations, RefCM can enable the discovery of new cell types, states, and relationships in single-cell transcriptomic data.

## 1 Introduction

Single-cell RNA sequencing (scRNA-seq) has opened new frontiers in biological research by capturing gene expression profiles at individual cell resolution [1–5]. This technology reveals the remarkable diversity within tissues, where populations previously thought to be homogeneous are now known to contain distinct cellular subtypes and states [6, 7]. A fundamental challenge in scRNA-seq analysis is cell type annotation: assigning biologically meaningful identities to cells based on their transcriptional signatures. Robust cell type identification underpins key biological investigations, from decoding developmental programs to elucidating disease mechanisms [8–10].

Traditional approaches to cell type annotation rely heavily on expert analysis of gene expression patterns. The standard pipeline involves clustering cells and identifying marker genes that distinguish each cluster [11]. These markers are detected through statistical tests for differential expression and interpreted using gene set enrichment (GSEA) analysis to match clusters with known cell type signatures [12]. While this approach has enabled numerous biological discoveries, it faces significant scalability challenges as datasets grow to hundreds of thousands of cells [13]. Manual annotation becomes increasingly impractical, requiring substantial time and expertise to resolve ambiguous cases and maintain consistency across studies[14–20]. The variability in marker gene selection and interpretation further complicates reliable annotation at scale [21].

The rapid growth of single-cell reference atlases [22, 23] has enabled a new paradigm for cell type annotation: *reference mapping*. This approach promises to automate and standardize annotation by leveraging existing expert-annotated datasets [24]. Reference mapping algorithms align query and reference data for supervised label transfer, often via a shared embedding space [25, 26]. A number of methods have been developed to tackle this problem: Seurat [27] uses dataset alignment through “anchors” followed by label transfer via weighted voting, scANVI [28] employs probabilistic modeling to learn cell embeddings and identities jointly, and CellTypist [29] applies logistic regression to harmonized data. Other widely used approaches include batch-integration methods such as SCALEX [30] and similarity-based annotation methods such as SingleR [31] and scmap [32]. However, these methods often struggle with biological and technical variations between reference and query data, particularly when annotating cells across different experimental conditions, tissues, or species [21, 33].

While cell-to-cell mapping approaches offer high resolution, they can be computationally intensive and sensitive to technical noise. This motivates a different strategy based on two key biological principles: cell types represent stable transcriptional states [34–37], and relative gene expression patterns are largely preserved across technologies [38–40]. Under these principles, we can transfer reference labels at the cluster level rather than at the level of individual cells, an approach we term *reference cluster mapping*. This coarser-grained strategy promises to be more robust and computationally efficient, yet has received comparatively less attention. Current methods like CIPR [41] and ClustifyR [42] rely primarily on correlation between averaged expression profiles, discarding valuable information about expression heterogeneity within clusters. This oversimplification particularly impacts performance in complex scenarios such as partial matching or hierarchical cell type relationships.

In this work, we present RefCM, a novel algorithm that addresses these challenges through a distinct approach to reference cluster mapping. By employing the Wasserstein metric from optimal transport (OT) theory, RefCM quantifies transcriptomic similarity between cell populations while preserving their internal heterogeneity. We frame annotation transfer as a graph matching problem solved via integer programming, enabling flexible matching constraints that can handle partial correspondences and hierarchical relationships. Our method demonstrates strong performance across diverse experimental scenarios, achieving consistently high accuracy in challenging cross-species comparisons where prior approaches often degrade. Moreover, the algorithm maintains this performance across different technologies and phenotypes while supporting scalable, practical runtimes on large atlases and variations in label resolution. RefCM improves upon existing approaches, providing more reliable automated annotation of single-cell data, particularly for complex comparative studies between species.

## 2 Results

### 2.1 The RefCM model

RefCM is an automated cell type annotation algorithm that leverages optimal transport theory to map query clusters to reference cell types. At its core, the algorithm compares transcriptional profiles between unannotated query cells and annotated reference cells to transfer cell type labels based on expression similarities. Formally, RefCM takes as input two gene expression matrices: a query dataset 𝒬 and a reference dataset ℛ. Additionally, it utilizes two mapping functions: *φ*, which assigns query cells to their cluster labels 𝒞, and *γ*_*r*_, which assigns reference cells to their known cell type annotations 𝒜_*r*_ — effectively defining a clustering of ℛ. The goal is to produce a mapping *γ*_*q*_ that assigns each query cell to either a reference cell type in 𝒜_*r*_ or marks it as a novel cell population, denoted by *θ*.

The RefCM workflow consists of three main steps. First, both datasets are projected into a shared embedding space to enable direct comparison. Second, RefCM computes a cost matrix 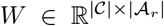 between query clusters and reference cell types using the Wasserstein metric. This metric captures the full distribution of gene expression within clusters, with lower values in *W* indicating lower transport costs, equivalently higher similarity, between populations. The cost matrix naturally represents a bipartite graph matching problem between query clusters and reference cell types, which RefCM solves through integer programming to handle differences in annotation resolution between datasets. The resulting matches provide labels for query clusters, with unmatched clusters designated as novel cell types *θ*. For a complete mathematical description of the algorithm, see Section 4.1.

### 2.2 Model evaluation

We conducted comprehensive experiments to evaluate RefCM against current state-of-the-art methods: Seurat [27], scANVI [28], CellTypist [29], CIPR [41], SingleR [31], scmap [32], clustifyr [42], SCALEX [30], SATURN [43], and a linear SVM classifier [21]. Our evaluation framework addresses three key types of annotation challenges: biological variation spanning both phenotypic differences and evolutionary distance between species, technical variation across sequencing technologies, and differences in annotation granularity between reference atlases including cell type hierarchies and resolution mismatches. We first evaluate performance on cross-technology, cross-phenotype, and cross-species annotation tasks. We then examine annotation granularity across different reference atlases, and finally assess runtime scaling across increasing dataset sizes.

To assess cross-technology annotation, we leveraged the scIB pancreas benchmark [44], which comprises 9 pancreas datasets collected using different sequencing technologies. We additionally evaluated cross-technology transfer on the PBMC Bench1 dataset [21] and on CellBench [45], a controlled cross-platform benchmark. For cross-phenotype comparison, we examined the impact of aging on cell type profiles using monkey adrenal gland datasets from [46]. We also evaluated cross-individual generalization using the Tabula Muris Senis dataset [47]. Finally, we evaluated cross-species annotation at different evolutionary distances. We began with the Allen Brain Atlas datasets [48, 49] to compare annotations between functionally distinct brain regions: the anterior lateral motor cortex (ALM) and primary visual cortex (VISp) from mouse brain samples, and the middle temporal gyrus (MTG) from human brain tissue. We then extended our analysis to more distant evolutionary relationships using frog and zebrafish embryogenesis datasets [50], where reduced gene homology presents additional challenges. For detailed descriptions of all datasets, see Section 4.6.

To enable fair comparison between methods, we adapted both RefCM and competing approaches to work across different annotation frameworks. Since many competing methods are *reference mapping* methods that operate on individual cells rather than clusters, we augmented them with majority voting (prefixed with `MV-`) to effectively convert them to *reference cluster mapping* methods. For cluster-level evaluations, we used the benchmark-provided oracle partitioning of query cells as an upper bound on reference cluster mapping performance [51], consistent with common annotation workflows in which cell types are first defined at the cluster level before assigning labels. Cell annotation accuracy was computed as closed-set accuracy: the percentage of query cells whose true type is present in the reference that were assigned the correct label, to avoid confounding comparisons by method-specific rejection/novelty detection procedures and sensitive threshold hyperparameters that are not uniformly available across approaches. See Section 4.7 for detailed benchmarking procedures.

### 2.3 RefCM performs robustly across technical and biological variation

Comprehensive benchmark evaluations show that RefCM consistently outperforms existing methods across diverse annotation challenges (Fig. 2). RefCM attains near-perfect accuracy across cross-technology tasks (scIB pancreas and PBMC Bench1), phenotypically varied monkey adrenal gland samples, and evolutionarily distant brain region comparisons, achieving perfect accuracy in all settings except Tabula Muris Senis and frog–zebrafish embryogenesis (where it remains the top-performing method). RefCM also achieves the best overall performance on Tabula Muris Senis group-*k*-fold cross-individual prediction, highlighting robust transfer across individuals. RefCM maintains high accuracy even for rare cell types on Tabula Muris Senis, correctly labeling clusters with fewer than 100 cells in the query or reference (Supplementary Fig. 1). Gains are most pronounced in cross-species brain annotations, where competing methods frequently fall below 65% accuracy even when aided by clustering information. These results highlight RefCM’s unique capability to overcome the technical and biological barriers that typically hinder accurate cell type annotation across diverse datasets.

**Fig. 1:**
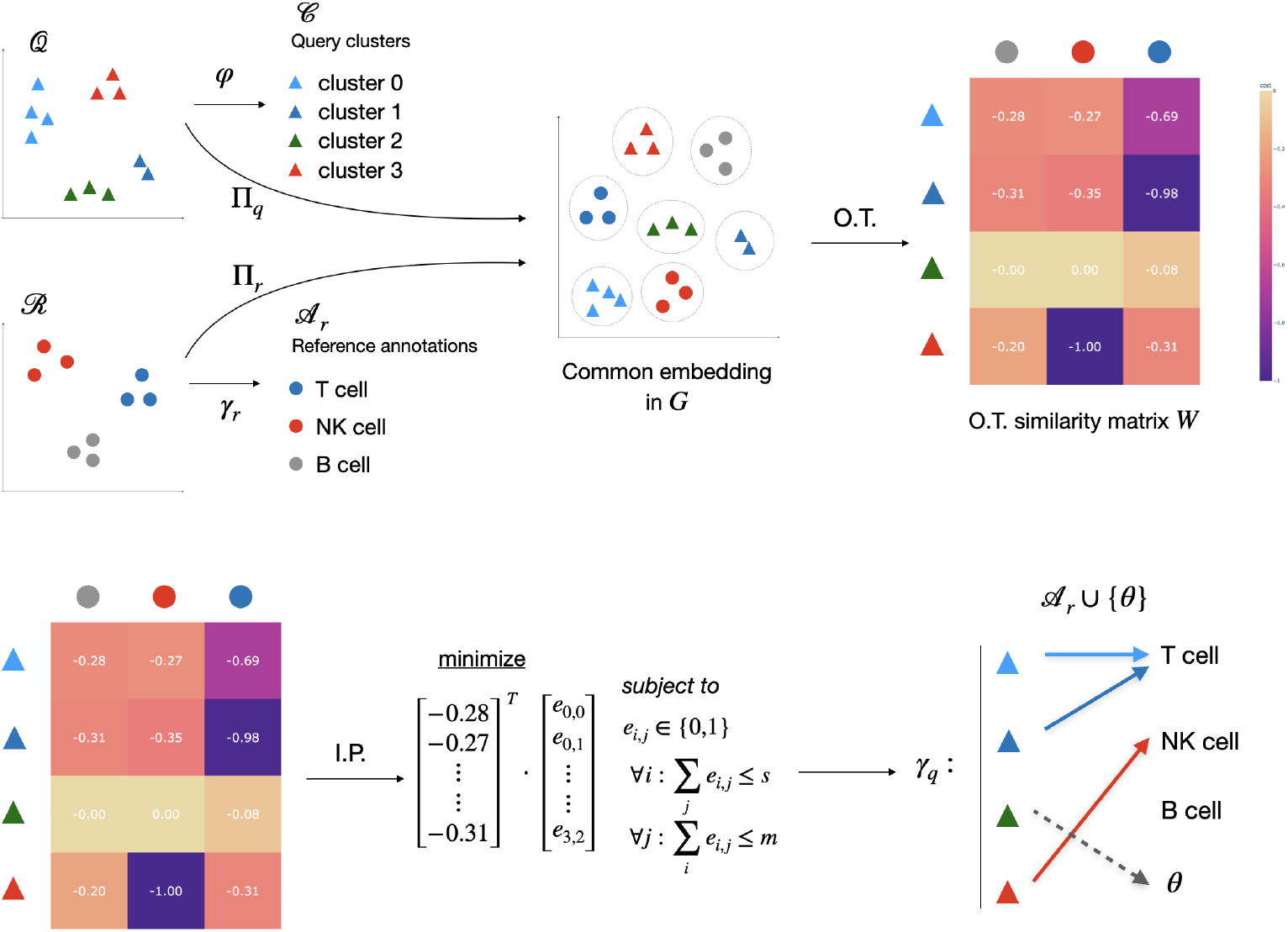
Overview of RefCM. Query and reference datasets are embedded in a common space (top-middle), where pairwise Wasserstein costs between clusters are computed (top-right). An integer program then determines the best cluster matches under given constraints (bottom-middle), which are used to annotate query clusters with reference labels. Unmatched clusters are labeled as novel (bottom-right).

**Fig. 2:**
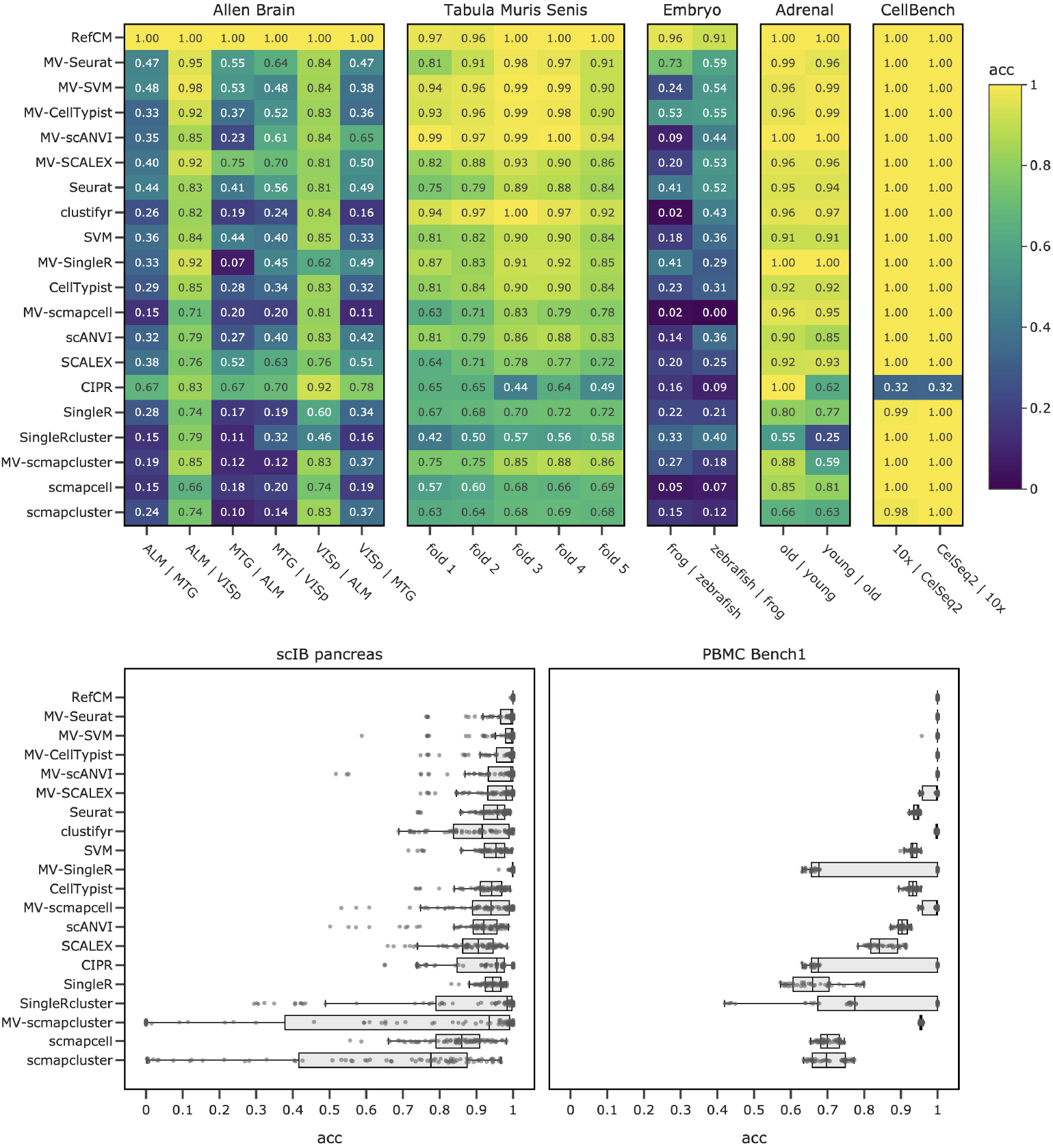
Benchmark performance of RefCM and baseline methods across diverse annotation scenarios. Methods are ordered by median accuracy across all tasks. Top row: accuracy heatmaps for Allen Brain cross-tissue and cross-species annotations, Tabula Muris Senis (droplet) group-*k*-fold cross-individual prediction, frog–zebrafish embryogenesis cross-species transfer, monkey adrenal gland cross-age transfer, and CellBench cross-technology evaluation. Bottom row: accuracy distributions for pairwise cross-technology annotations on scIB pancreas and PBMC Bench1 datasets. Throughout, the notation *X* | *Y* denotes annotation of query dataset *X* given *Y* as reference.

### 2.4 Accurate mapping across distant evolutionary relationships

To demonstrate RefCM’s effectiveness in cross-species annotation, we focused on challenging cases with significant evolutionary distance. Fig. 3 visualizes performance differences between methods when annotating human MTG using mouse ALM as reference. While competing methods show considerable misclassification (red cells), RefCM maintains high accuracy across the diverse cell populations present in these evolutionarily distinct brain regions.

**Fig. 3:**
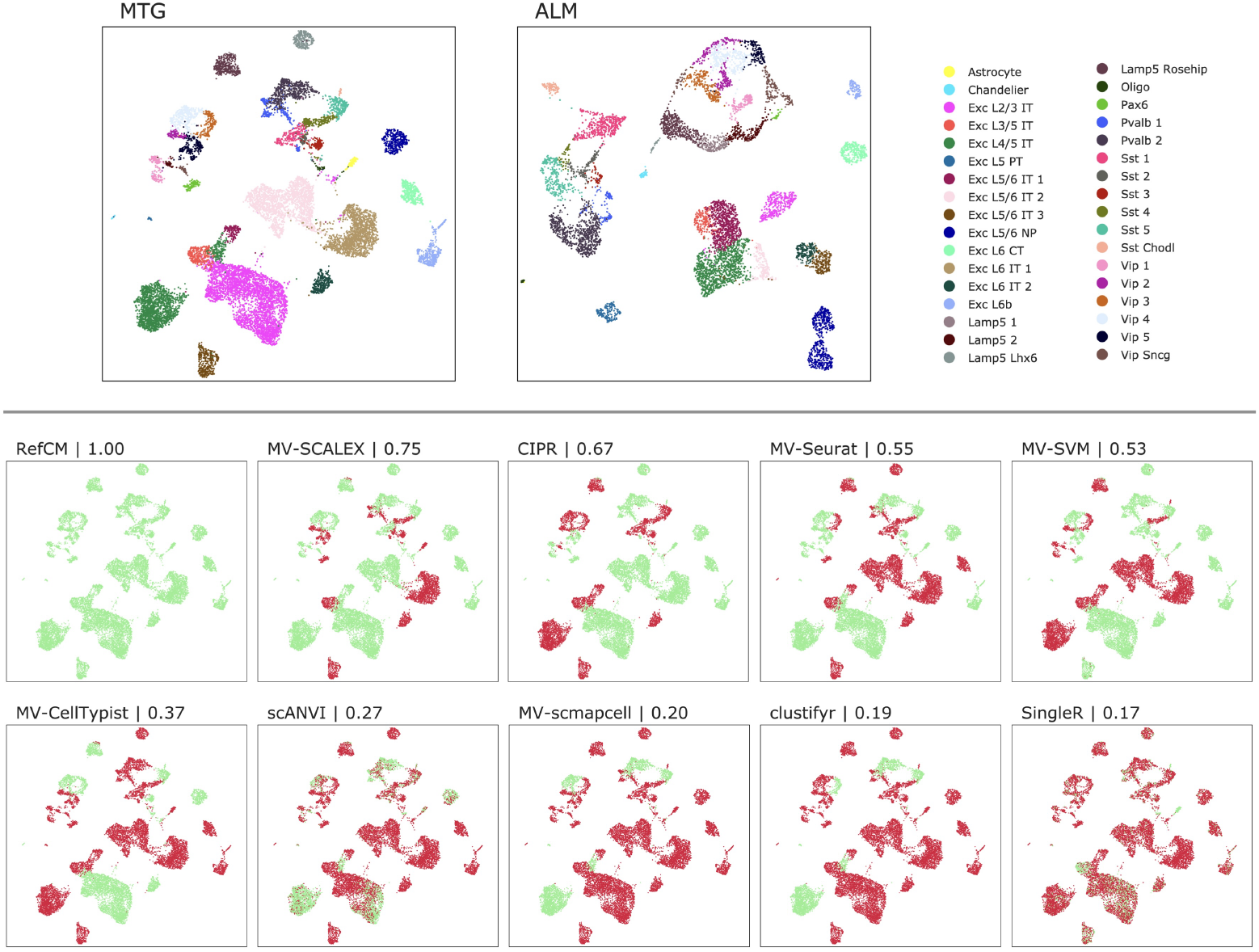
Cross-species annotation of MTG (human) given ALM (mouse) as reference. UMAP plots (top). Correct (green) and incorrect (red) cell annotations with accuracies for each method (bottom). For each method, we show the best-performing variant on this task (e.g., `SingleR`, `SingleRcluster`, or `MV-SingleR`).

On the evolutionarily distant frog–zebrafish embryogenesis transfer task, RefCM performs on par with SATURN [43] when the latter is augmented with majority voting to leverage clustering information; both methods correctly map 25 out of 28 common cell types. Notably, RefCM’s explicit handling of novel cell types enables it to identify 5 out of 14 non-common cell types, providing an explicit treatment of novel populations rather than forcing a best-match assignment. In contrast, several cell-level annotation baselines are more sensitive to distribution shift between reference and query data in this setting, which we observe as increased misclassification and reduced robustness under cross-species transfer.

### 2.5 Resolving resolution differences and annotation hierarchies

To evaluate RefCM’s ability to handle differences in annotation resolution and hierarchical relationships, we leveraged the Allen Brain dataset’s multi-level annotation structure. These datasets are particularly well-suited for this analysis as they combine cross-species variation with two distinct levels of annotation granularity: a coarse level with 3 super-types and a fine-grained level with 34 cell types (Fig. 4). By allowing many-to-one and one-to-many mappings (i.e., permitting cluster merging and splitting), we tested RefCM’s performance on all six possible dataset combinations, considering both coarse-to-fine and fine-to-coarse annotation scenarios. RefCM successfully recovered the hierarchical relationships in all cases, accurately mapping super-types to their constituent subtypes and vice versa. This ability to transfer labels across mismatched resolutions while maintaining accurate cross-species mapping is not supported by many existing approaches, which typically assume comparable annotation granularity between reference and query datasets.

**Fig. 4:**
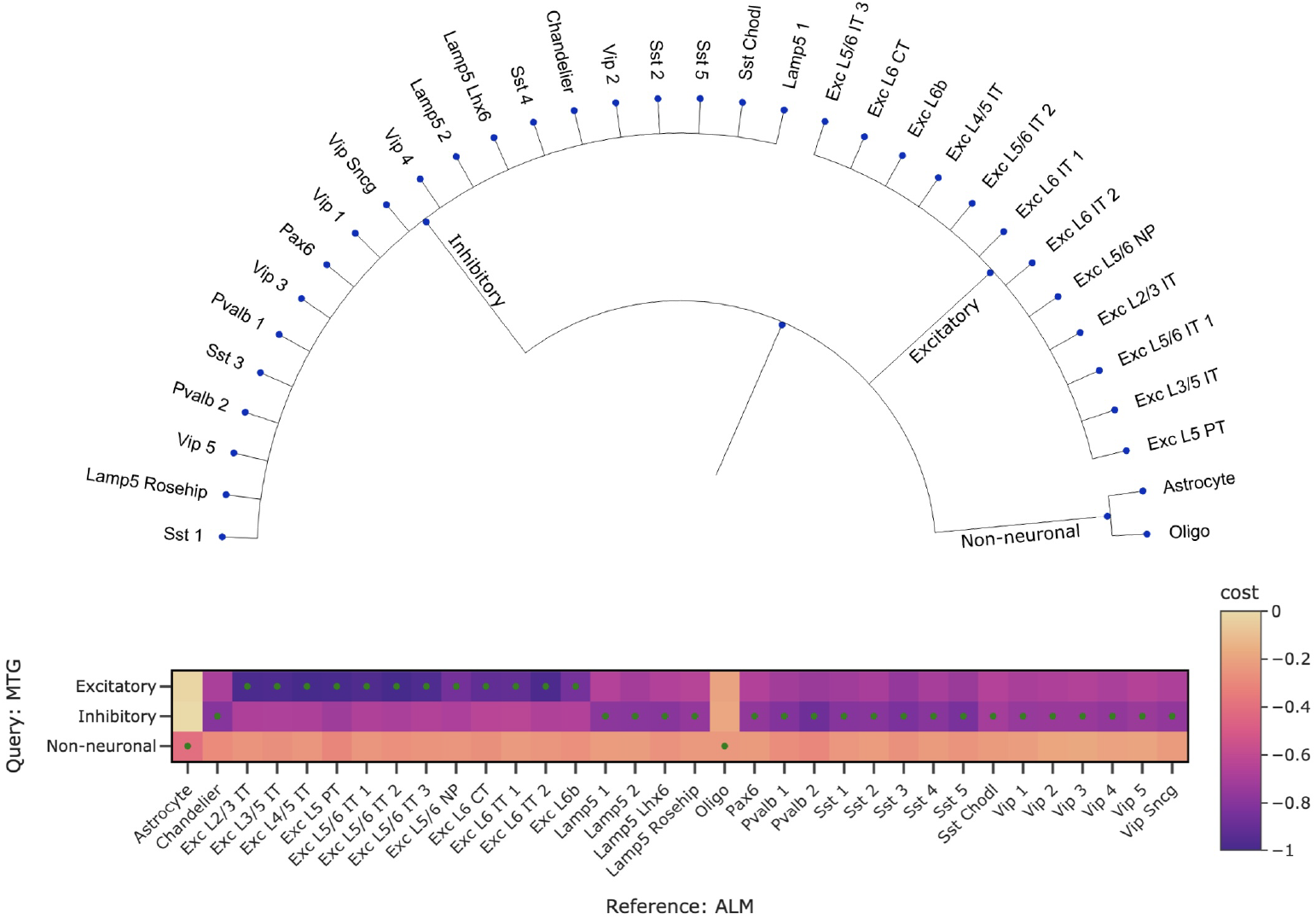
Top: Allen Brain Atlas cell type hierarchy. Bottom: Example of mapping coarse type-level labels from VISp to the granular level on the ALM dataset. The color gradient illustrates the Wasserstein distances computed between clusters in the OT-step, and dots indicate the final mappings returned by RefCM.

### 2.6 Runtime scaling and atlas-scale deployment

Beyond accuracy, practical atlas-scale annotation requires methods that remain tractable as the number of cells grows and that support repeated query annotation. Single-cell annotation methods differ substantially in both their compute profile and how costs split between reference setup and query-time prediction. Many approaches are primarily CPU-bound and benefit from multicore parallelism, whereas neuralnetwork methods typically achieve their best throughput with GPU acceleration. Separately, several methods explicitly separate a costly reference training step from lightweight inference (e.g., CellTypist and SVM), enabling a model trained once on a large atlas to be reused across many downstream queries.

We benchmarked wall-time runtimes on Tabula Muris subsamples across *n* = 5,000–200,000 cells (80/20 reference/query split; 10–100 cell types), reporting mean runtime over 5 runs on an NVIDIA GH200 system (Fig. 5). RefCM scales favorably across all tested sizes and completes end-to-end annotation in 151 seconds at *n* = 200,000, comparable to Seurat (146 seconds) and substantially faster than GPU-accelerated baselines even when run with dedicated GPU resources in our benchmark (SCALEX: 3407 seconds; scANVI: 4485 seconds), as well as several widely used methods (CIPR: 1428 seconds; scmapcell: 1173 seconds; SingleR: 2957 seconds). These results indicate that RefCM achieves strong accuracy while remaining practical at atlas scale without relying on GPU acceleration.

**Fig. 5:**
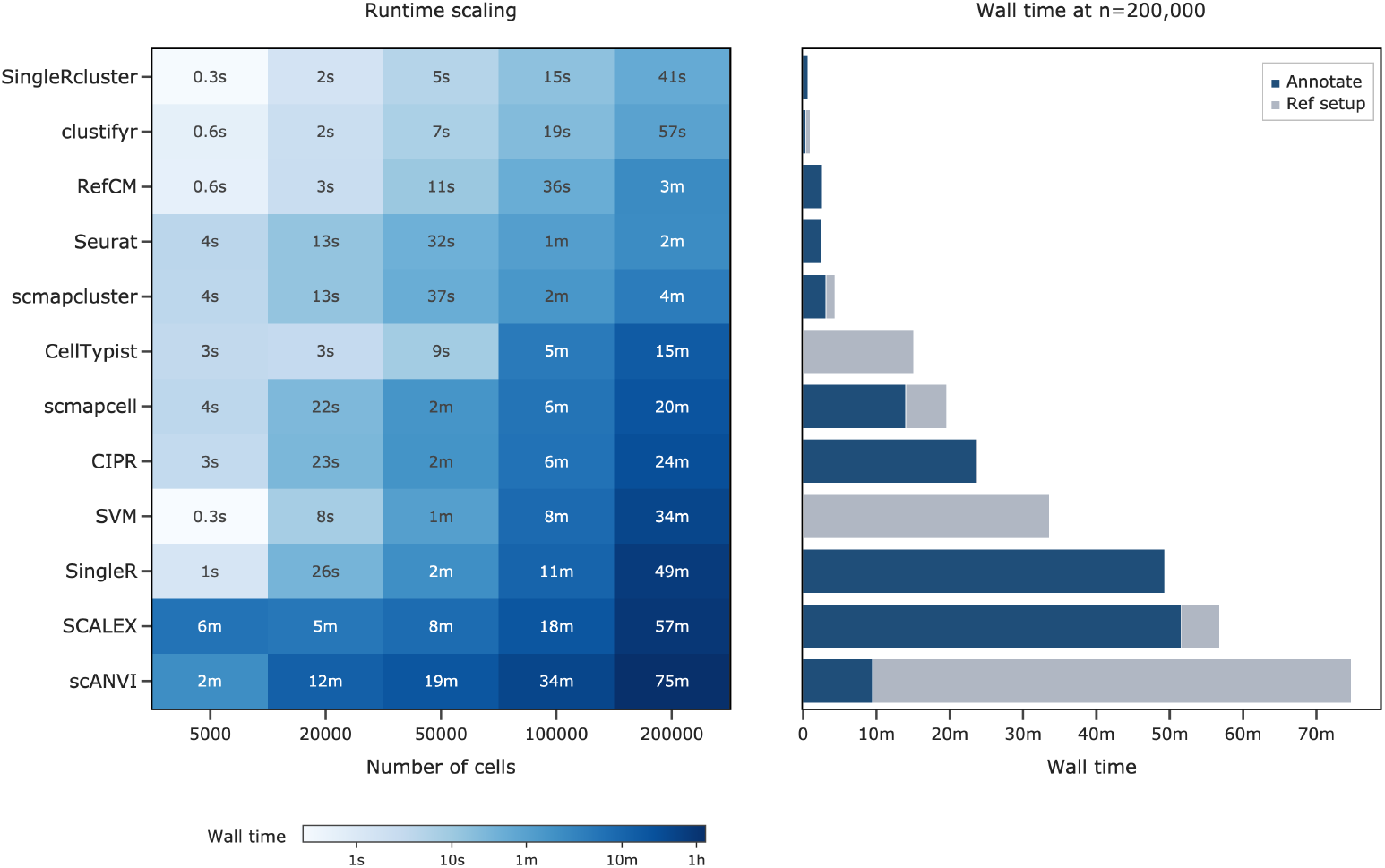
Wall-time runtimes (mean over 5 runs) on Tabula Muris subsamples with fixed sizes *n* ∈ *{*5 000, 20 000, 50 000, 100 000, 200 000} (80/20 reference/query split) and corresponding increasing numbers of cell types (10–100). Left: runtime heatmap across methods and dataset sizes (log scale); methods are sorted by mean wall time across sizes. Right: runtime decomposition at *n* = 200,000 into reference setup (training where applicable) and query annotation. Experiments were run on an NVIDIA GH200 system.

The runtime decomposition at *n* = 200,000 further clarifies deployment tradeoffs. CellTypist and SVM devote nearly all runtime to reference training (904 and 2015 seconds, respectively) but then achieve sub-second query inference (0.57 and 0.34 seconds), which is advantageous when a fixed pretrained classifier can be reused. In contrast, methods dominated by query-time computation (e.g., CIPR with 1417 seconds spent in annotation) incur comparable cost whenever the query changes. RefCM is CPU-bound and parallelizes naturally across query–reference cluster pairs in the OT step; the final integer-program step is negligible at the cell-type counts considered here. While RefCM does not provide a standalone pretrained classifier, it offers fast end-to-end runtimes together with strong performance, making it well suited to iterative analysis settings where references and queries vary across runs.

## 3 Discussion

RefCM introduces a novel approach to automated cell type annotation by combining optimal transport theory with integer programming. At its core, the method quantifies transcriptomic similarity between reference and query clusters through optimal transport, translating the annotation problem into a flexible graph matching framework. This formulation naturally supports partial correspondences, including novel populations, and transfer across mismatched annotation resolutions, addressing key challenges in single-cell annotation.

We demonstrated RefCM’s strong performance across diverse experimental settings, from technical variation across sequencing technologies to biological differences in age and species. The method particularly excels in cross-species annotation, where distribution shifts pose significant challenges for existing approaches. The frog– zebrafish task is especially illustrative: despite limited gene overlap, and the fact that homologous genes can diverge in function across species, RefCM performs remarkably well. SATURN addresses this setting by leveraging protein embeddings, yet RefCM achieves comparable (and slightly better) results using only the intersection of highly variable genes as detailed in the methods section. This suggests that optimal-transport distances between expression distributions can extract substantial cross-species signal even under imperfect homology, positioning RefCM as a competitive alternative in settings where embedding-based approaches are typically used. These results also suggest the potential for future work to integrate protein embeddings with RefCM for further gains.

In addition to accuracy, our runtime benchmarks indicate that RefCM remains practical at atlas scale. RefCM is CPU-bound and benefits directly from multicore parallelism, since query–reference cluster pairs can be evaluated independently in the OT step, while the final integer-program mapping is negligible compared to the OT computation. In our benchmark at *n* = 200,000, RefCM achieves competitive end-to-end runtimes without requiring GPU acceleration, whereas many neural-network baselines effectively require GPU acceleration to remain computationally tractable at this scale. This makes RefCM a practical choice for day-to-day single-cell workflows, where analyses are refined iteratively and runtime and compute costs can be limiting. There are a number of future directions to explore. The success of RefCM suggests that the notion of cell type is largely captured by the empirical distribution of expression vectors. Nonetheless, a natural question is whether improved representations can provide more refined cell type information, for example by defining the OT cost in learned embedding spaces (e.g., scVI- or SATURN-style embeddings) in addition to gene space. More generally, we might search for a refined notion of cell type that captures subtler biological phenomena than are accessible from expression alone. In another direction, since the effectiveness of RefCM hinges on the accurate clustering of the query dataset, an important next step is to develop clustering procedures that are explicitly informed by the reference. Current approaches typically fall into two categories: unsupervised clustering or supervised classification. While unsupervised clustering can uncover novel populations, its outcomes are sensitive to hyperparameter choices (e.g., the number of clusters or resolution) that are difficult to set in advance. Supervised classification methods can leverage reference information, but are not designed to discover new clusters or populations, and most do not explicitly incorporate the local geometry and structure of the query object they perform inference on. A promising middle ground is a supervised (or joint) clustering approach that can transfer structure from one or more references while retaining the ability to identify novel populations. Finally, RefCM currently relies primarily on cross-dataset distances between clusters. It would be interesting to incorporate within-dataset geometry, for example by using OT to define graphs of cluster–cluster relationships within the query and reference and regularizing the integer program to preserve these structures, in a manner related in spirit to Gromov–Wasserstein formulations.

## 4 Methods

### 4.1 Algorithm Overview

1. **Input:** RefCM takes two primary data inputs.
  - A query dataset 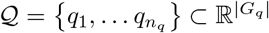 containing gene expression vectors for *n*_*q*_ cells, usually count and log-normalized expression values, where each cell’s expression is measured across a set of genes *G*_*q*_.
  - A reference dataset 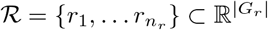containing gene expression vectors for *n*_*r*_ cells, measured across a set of genes *G*_*r*_. *Note*: The gene labels *G*_*q*_ and *G*_*r*_ may differ, which is particularly common when comparing across species. RefCM can also work with data that has already been embedded in a common space ℰ .
2. **Input:** The reference dataset ℛ includes cell type labels:
  - A set of cell types 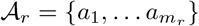
  - An annotation map *γ*_*r*_ : ℛ ↦ 𝒜_*r*_ that assigns each reference cell to one of these types, i.e. ∀*r* ∈ ℛ, *γ*_*r*_(*r*) ∈ 𝒜_*r*_. These annotations effectively partition the reference cells into clusters 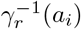, where each group contains all cells of a particular type.
3. **Problem:** Our task is now finding the set of cell types 𝒜_*q*_ ⊆ 𝒜 _*r*_ *θ* that our query dataset presents, where *θ* denotes a novel cell-type — or rather one that is not present in our reference — and a corresponding assignment *γ*_*q*_ : 𝒬 ↦ 𝒜_*q*_ such that *γ*_*q*_(*q*) is the correct type annotation for every cell *q* ∈ 𝒬.
4. **Problem:** We make the further assumption that the query 𝒬 is equipped with a clustering (i.e., partition) 𝒬 = ∪_*i*_𝒞_*i*_, and we make the simplifying assumption that:

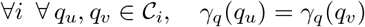

That is, *γ*_*q*_ can be regarded as an assignment {𝒞 _*i*_} *→ 𝒜*_*q*_. Note that we do not require this assignment to be injective; in some cases multiple clusters map to the same type, which we will call “merging”. In the “splitting case”, a single cluster may map to multiple reference types; then the cluster-level annotation becomes set-valued, e.g.,

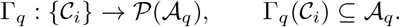
5. **Algorithm:** We embed the query and reference data set in a common space ℝ^| 𝒢|^. For example, a particularly simple embedding uses only the set of genes common to both reference and query, *G*_*q*_ ∩ *G*_*r*_.
6. **Algorithm:** Next, we compute the Wasserstein optimal transport cost from mapping the clusters of the query dataset to the clusters from the annotated reference dataset. Specifically, we regard a cluster as specifying the empirical distribution on its constituent cells and compute the Wasserstein loss between these distributions to compare clusters. This results in a cost matrix *W* between query and reference clusters in the embedding space that captures the similarity of the underlying population distributions [52].
7. **Algorithm:** The resulting matrix defines a bipartite graph matching problem, which we solve using integer programming with additional biological constraints. The parameters and constraints are described in more detail in Section 4.4.
8. **Output:** Based on the optimal matching, we assign reference cell type labels to query clusters. Query clusters without matches are labeled as novel cell populations (*θ*).

### 4.2 Joint Embeddings

To accurately compare genetic expression variability between query and reference cell types, it is essential to first map both datasets to a joint embedding space. Here, we begin by log-normalizing both query and reference datasets and select highly variable genes (HVG) from each to reduce the original gene expression spaces to subspaces that preserve relevant biological information [5, 27, 53]. By performing HVG selection independently on both query and reference datasets, we identify the most informative genes in each context. We then take the union of these high-variable gene sets while ensuring that only common genes between the query and reference datasets are retained. This unified set of high-variable genes forms the basis of the joint embedding space, enabling cellular comparisons in the following OT step. More formally, if 𝒬 and ℛ are our query and reference datasets, respectively, and *G*_*q*_ and *G*_*r*_ represent their gene sets, our embedding space is then 𝒢 = (`hvg`(𝒬) ∩ *G*_*r*_) ∪ (`hvg`(ℛ) ∩ *G*_*q*_). Our embedding maps *π* _𝒬_ and *π* _ℛ_ are then simply the canonical projections of 𝒬 and ℛ into this common subspace.

### 4.3 Optimal Transport & Wasserstein distance

Optimal transport (OT) is a mathematical framework for comparing probability distributions by reallocating mass (or probability) from one distribution to another in the most cost-effective manner, where the cost is defined in terms of a distance metric between points in the distributions. Unlike simpler metrics that compare summary statistics (e.g., means or variances), OT considers the full shape of distributions, making it particularly valuable for comparing heterogeneous cell populations.

Consider two discrete distributions: a source distribution *µ* with points *{x*_*i*_*}* and a target distribution *ν* with points {*y*_*j*_} . The goal of optimal transport is to find a transport plan *π* that reallocates the mass from *µ* to *ν* while minimizing the total transportation cost:

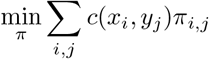

subject to the constraints:

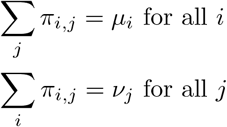

where *π*_*i,j*_ represents the amount of mass transported from *x*_*i*_ to *y*_*j*_, *µ*_*i*_ and *ν*_*j*_ are the masses at points *x*_*i*_ and *y*_*j*_, and *c*(*x*_*i*_, *y*_*j*_) is the ground-cost (transport cost) of moving a unit of mass between these points.

The minimal transportation cost is known as the Wasserstein distance (or Earth Mover’s distance), which quantifies the similarity between distributions. Intuitively, this can be understood as the minimum effort required to reshape one distribution into another.

In the RefCM algorithm, we apply this concept to compare cell type distributions between query and reference datasets (Figure 6). We treat each cluster as a discrete uniform distribution over its constituent cells, with the transportation cost between cells defined by a ground cost in the shared gene expression space (typically Euclidean distance or a cost derived from the inner product between embeddings). The Wasserstein distance quantifies the cost of transforming the distribution of gene expressions in the query cluster into the distribution in the reference dataset’s cluster, capturing the structural differences between them (with lower cost for more similar populations). This cost metric forms the basis for further computation to optimally map query and reference cell types between datasets.

**Fig. 6:**
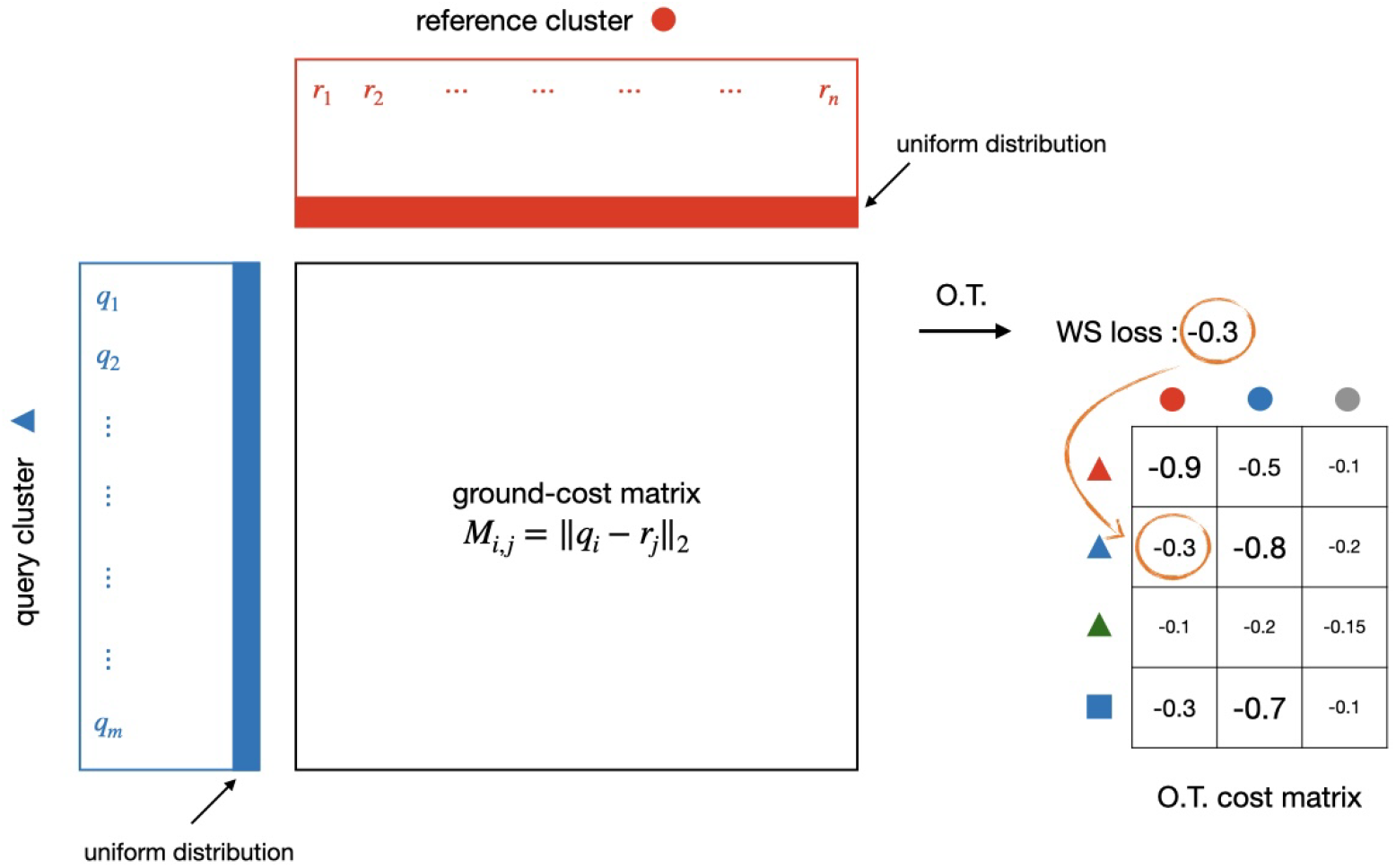
Optimal Transport between query and reference clusters.

To construct the cost matrix used in the next step, we compute the pairwise Wasserstein distance between each query cluster 𝒞_*i*_ and reference cell type 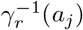. Let *c*(·, ·) denote the ground-cost function in the shared embedding space (e.g. Euclidean distance). We define the unnormalized Wasserstein cost matrix

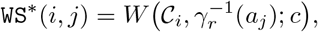

where *W* (·,· ; *c*) denotes the Wasserstein (Earth Mover’s) distance between the two clusters under the ground cost *c*, with uniform weights over cells in each cluster. We then rescale these values to the interval [−1, 0] via

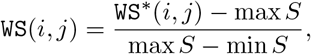

where *S* = {`WS`^∗^(*i, j*) | 1 ≤ *i ≤ m*_*q*_, 1 ≤ *j ≤ m*_*r*_ }. The resulting matrix `WS` is therefore a normalized Wasserstein cost matrix, with lower values indicating lower transport cost (i.e. greater similarity) between populations.

### 4.4 Integer programming for relaxed graph matching

To determine the optimal cluster-to-cluster mappings between query and reference datasets, we devised an integer program based on the Wasserstein cost (WS) matrix obtained in the previous step. This cost matrix represents a complete bipartite graph, where clusters (or cell types) from the query and reference datasets form the graph’s nodes, and the edge weights correspond to the WS cost values. The integer programming framework enables us to identify the set of edges that minimizes the total mapping cost while adhering to a set of prescribed constraints. This approach effectively disentangles the optimal set of cluster mappings, ensuring accurate and efficient alignment of cell types between the query and reference datasets. We now describe the matching constraints, which can be adjusted depending on the application.

Formally, we define *G*(*V, E*) as a fully connected bipartite graph where edges *e*_*i,j*_ connect query cluster *i* to reference cell type *j*, with edge weight:

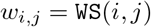

For each edge in the graph, we introduce a binary decision variable *x*_*i,j*_ indicating whether that edge is included in the optimal mapping. Our objective is to minimize the sum of costs over the chosen edges

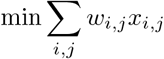

subject to the constraints:

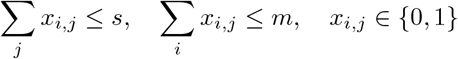

Due to differences in clustering scales, granularities, or cell distributions, it is often necessary to match multiple clusters from the query set to a single reference cluster, a phenomenon we term “cluster merging.” Conversely, a single cluster from the query set may correspond to multiple reference clusters, which we refer to as “cluster splitting.” This consideration is further motivated by the hierarchical structure of cell types, where we may need to match finer-grained clusters to a broader parent type or, alternatively, more coarse clusters to specific reference subtypes.

The first constraint ensures that each cluster in the query set can be connected to up to *s* ∈ ℕ clusters in the reference dataset, allowing for cluster splitting into as many as *s* reference clusters. Conversely, the second constraint guarantees that each cluster in the reference set can be mapped to up to *m* ∈ ℕ clusters in the query set, permitting the merging of up to *m* query clusters into a single reference cluster. The third constraint restricts the linear program decision variables to binary values, ensuring the optimal decision variable outcomes align with the intended matching framework.

Finally, because the reference dataset may not encompass all cell types present in the query dataset, it is essential to account for non-common cell types and facilitate cell type discovery. To address this, we establish a threshold for identifying novel cell types by discarding high-cost query–reference correspondences. Specifically, we set all costs *w*_*i,j*_ whose normalized value lies in the worst *p* fraction of the global cost distribution, i.e., above the (1 − *p*)-quantile *τ*_*p*_ of the normalized cost matrix, to infinity. We then threshold edges as

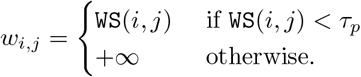

This ensures that the integer programming solver will never select these edges. By constraining all remaining costs to be between −1 and 0, the solver favors returning a set of edges that minimizes the overall cost while adhering to the constraints listed above. This approach effectively detects and handles non-common cell types, enabling accurate cell type discovery.

### 4.5 Implementation

RefCM is implemented in Python. We use POT [54] to compute Wasserstein distances and PuLP [55] to formulate the integer programs, solved with the GNU Linear Programming Kit (GLPK) [56]. Through POT, RefCM supports alternative OT formulations (e.g., Sinkhorn-regularized, unbalanced, and Gromov–Wasserstein OT), with EMD used by default. The optimal transport step dominates computation and is parallelized across query–reference cluster pairs using a `ThreadPoolExecutor`. The default EMD solve runs in POT’s compiled backend and releases the Python GIL during the core computation, enabling multicore parallelism without duplicating large in-memory objects. The subsequent integer-program step is negligible compared to OT. We report end-to-end runtime scaling and a breakdown of reference setup versus query annotation in Fig. 5. The RefCM package is available on PyPI, with source code and documentation on GitHub.

### 4.6 Datasets

Across all experiments, we use the cell-type annotations distributed with each dataset/benchmark release as ground truth, without re-clustering or otherwise reannotating cells. When a dataset provides a predefined cross-dataset label space, we evaluate in that label space (e.g., Allen Brain at the 3- and 34-population levels [48] [49]; frog–zebrafish using the SATURN type mapping [43]). For Tabula Muris Senis, we evaluate cross-individual generalization using group-*k*-fold splits by mouse identity to prevent individual-level leakage and mitigate instability from highly variable pairwise overlaps in cell-type composition across mice.

**Table 1:**
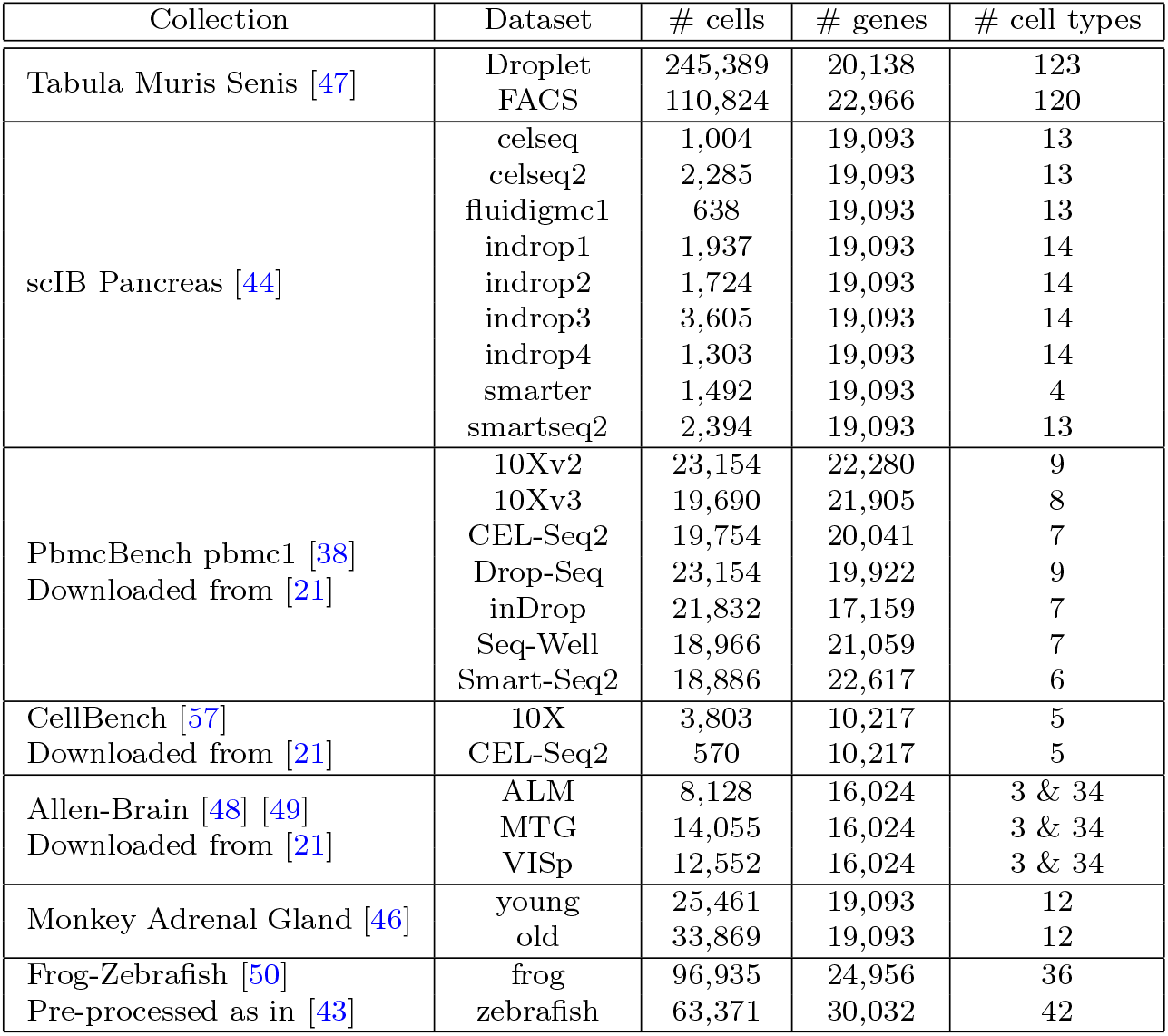
Summary of datasets used.

### 4.7 Performance benchmarking

We compared RefCM to widely used cell type annotation baselines and followed the default or recommended workflows in the corresponding official implementations. Unless otherwise noted, we report closed-set accuracy and disable method-specific rejection/novelty detection options to ensure comparability across approaches. We provided each method with its expected input representation: log-normalized expression for CellTypist [29], Seurat [27], CIPR [41], SingleR [31], scmap [32], and clustifyr [42], and raw counts for scANVI [28] and SCALEX [30].

All runtime experiments were run on a Lambda NVIDIA GH200 (Grace Hopper) instance, representative of the heterogeneous CPU+GPU nodes increasingly common in modern academic and institutional HPC environments. The system pairs a 72-core NVIDIA Grace CPU with 480 GB of LPDDR5X system memory (ECC) and an NVIDIA H100 (Hopper) Tensor Core GPU with 96 GB HBM3, connected by a cache-coherent NVLink-C2C interconnect (up to 900 GB/s). For R-based baselines, we evaluated common parallel execution configurations (including `future()`-based and `BiocParallel`-based backends where supported) and report runtimes for the fastest supported configuration for each method. For Python-based baselines, we likewise enabled method-supported parallelism (e.g., `n jobs`/`n workers` where exposed) and used the fastest supported configuration where applicable. Finally, we do not report memory usage: observed memory footprints depend strongly on the parallelization strategy (e.g., process-based workers vs. threads, object copying vs. shared memory, and garbage collection behavior), and for GPU-accelerated methods additionally on host–device staging and caching, making fair, comparable cross-method accounting difficult.

#### 4.7.1 RefCM

Across all benchmarks, we ran RefCM with fixed default parameters: Euclidean distance as the ground cost, discovery threshold 0.0, the Earth Mover’s Distance solver (`emd`) from POT, and the HVG embedding (Section 4.2). We set *s* = *m* = 1 for standard benchmarks to enforce one-to-one mappings and keep accuracy well-defined under splitting/merging, and set *s* = *m* = −1 only for hierarchy recovery experiments (Section 2.5). For HVG selection, we used the union of the top-1200 HVGs selected independently in the query and reference, yielding ∼2000 joint features in practice depending on overlap, comparable to the 2000-HVG convention used by baseline methods; our ablation (Supplementary Fig. 2) confirms that performance is stable across the 500–3000 range.

#### 4.7.2 CIPR

Following the recommended workflow and documentation, we used Seurat preprocessing to obtain `log1p`-normalized expression and computed query cluster marker statistics with `FindAllMarkers`. We provided CIPR with this differential-expression table and used the `logfc dot product` mode. For the reference, we constructed a custom reference profile by averaging expression within each reference label (`AverageExpression`) and supplied this as `custom reference`. Each query cluster was assigned to the reference label with the highest CIPR identity score.

#### 4.7.3 clustifyr

We ran clustifyr in its standard correlation-based cluster annotation mode. We constructed a reference profile matrix by averaging log-normalized expression within each reference label (`average clusters`) and computed query cluster profiles analogously from the provided query clustering. We then used `clustify` to assign each query cluster to the reference label with the highest correlation (Spearman correlation by default).

#### 4.7.4 SingleR

We used SingleR following the Bioconductor vignette. We constructed `SingleCellExperiment` objects for reference and query, computed log-normalized expression values, and ran SingleR using the benchmark-provided reference labels. We evaluated both the default per-cell mode and the cluster-aware mode via the `clusters=` option. In figures, we refer to the per-cell method as `SingleR` and the cluster-aware variant as `SingleRcluster`. Since `SingleRcluster` is not equivalent to majority-voting per-cell predictions, we also report a majority-vote aggregation of `SingleR` cell-level predictions.

#### 4.7.5 scmap

We ran scmap following the Bioconductor workflow. We used scmap’s feature selection (`selectFeatures`; 500 genes) on the reference and built either a cell-level index (`indexCell`) or a cluster-centroid index over reference labels (`indexCluster`). We then annotated each query cell with `scmapCell` or `scmapCluster`, respectively. For cluster-level evaluation, we aggregated both per-cell prediction methods by majority vote within each query cluster. We set scmap’s rejection threshold to 0 to avoid conflating unassigned predictions with misclassifications under closed-set accuracy. In figures, we refer to these as `scmapCell` and `scmapCluster`, respectively.

#### 4.7.6 Seurat

We ran Seurat label transfer using the standard anchor-based workflow. We normalized and `log1p`-transformed both datasets, selected variable features, computed anchors with `FindTransferAnchors` (`dims=1:30`), and transferred labels to the query using `TransferData` with default settings.

#### 4.7.7 CellTypist

Following the official documentation, we provided CellTypist with library-size normalized expression (scaled to 10,000 counts per cell) followed by `log1p`. We trained CellTypist on the reference labels using the reference HVGs (2000 genes). At prediction time, we restricted to the subset of these HVGs present in the query; if this gene set differed from the one used for training, we retrained CellTypist on the reference using the restricted gene set before annotating the query.

#### 4.7.8 scANVI

We ran scANVI (scvi-tools) [28] on raw UMI counts, restricting both reference and query to the top-2000 HVGs. We first trained a two-layer scVI model [58] on the reference (dropout 0.2) and then initialized scANVI from this model. We fine-tuned scANVI on the reference and subsequently adapted it using the query as in the standard scArches workflow, and used the resulting model to predict query cell types.

#### 4.7.9 SCALEX

We ran SCALEX following the official documentation. We provided raw count matrices and used SCALEX’s built-in preprocessing, which performs library-size normalization, `log1p` transformation, HVG selection (2000 genes by default), and scaling prior to model fitting. We trained SCALEX on the reference and projected the query into the learned latent space using the provided projection workflow. To avoid dropping cells used for evaluation, we disabled SCALEX’s default query-side QC filtering (cell filtering by `min features`). We then applied SCALEX’s label-transfer utility on the latent representations to obtain query predictions.

#### 4.7.10 SVM

We trained a linear SVM classifier (scikit-learn `LinearSVC`) [59] on the reference using the benchmark-provided labels and restricting to the reference HVGs (2000 genes). At prediction time, we restricted to the intersection of genes between reference and query; if the intersection differed from the training gene set, we retrained `LinearSVC` on the reference using the intersected genes before predicting query labels.

## Declarations

Some journals require declarations to be submitted in a standardized format. Please check the Instructions for Authors of the journal to which you are submitting to see if you need to complete this section. If yes, your manuscript must contain the following sections under the heading ‘Declarations’:

- Funding: N/A
- Conflict of interest/Competing interests (check journal-specific guidelines for which heading to use): N/A
- Ethics approval and consent to participate: N/A
- Consent for publication: N/A
- Data availability: All data is available on our [GitHub repository](https://github.com/vgalanti/RefCM)
- Materials availability: N/A
- Code availability: All code is available on our [GitHub repository](https://github.com/vgalanti/RefCM)
- Author contribution: N/A

If any of the sections are not relevant to your manuscript, please include the heading and write ‘Not applicable’ for that section.

