## Supplementary Information for "Automated Cell Type Annotation with Reference Cluster Mapping"

### Supplementary Figures

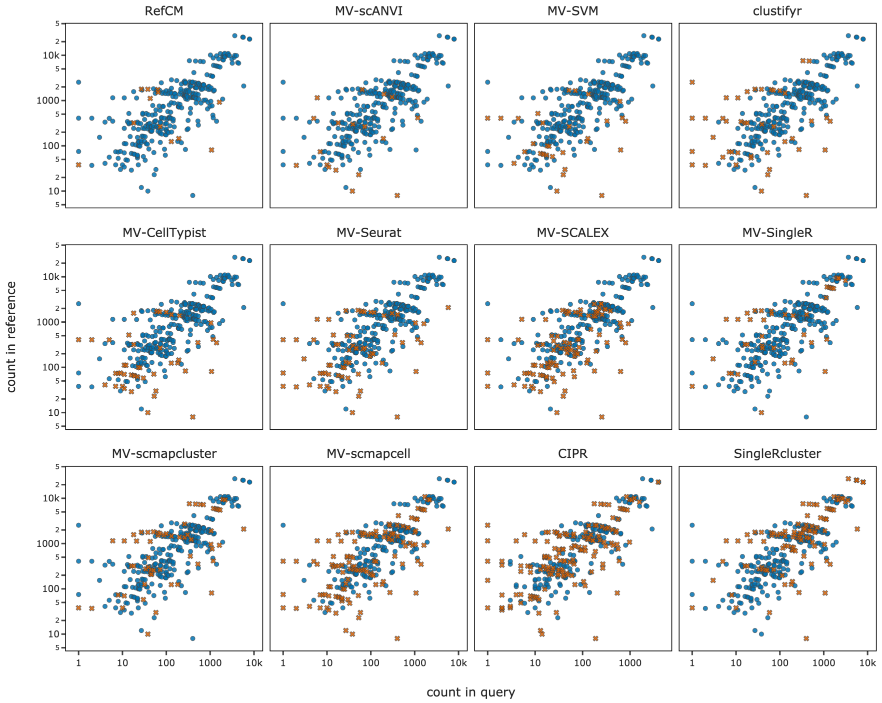


Supplementary Figure 1: Annotation accuracy vs. cell type abundance. Each point represents a cell type from the Tabula Muris Senis (droplet) benchmark, aggregated across group-$k$-fold cross-validation splits (grouped by mouse ID). Position indicates cell count in query (x-axis) and reference (y-axis); blue circles denote correctly annotated cell types, orange crosses denote incorrectly annotated. Low-abundance cell types in either query or reference tend to yield lower annotation accuracy, with methods differing in their robustness to class imbalance. Methods ordered by descending mean accuracy.

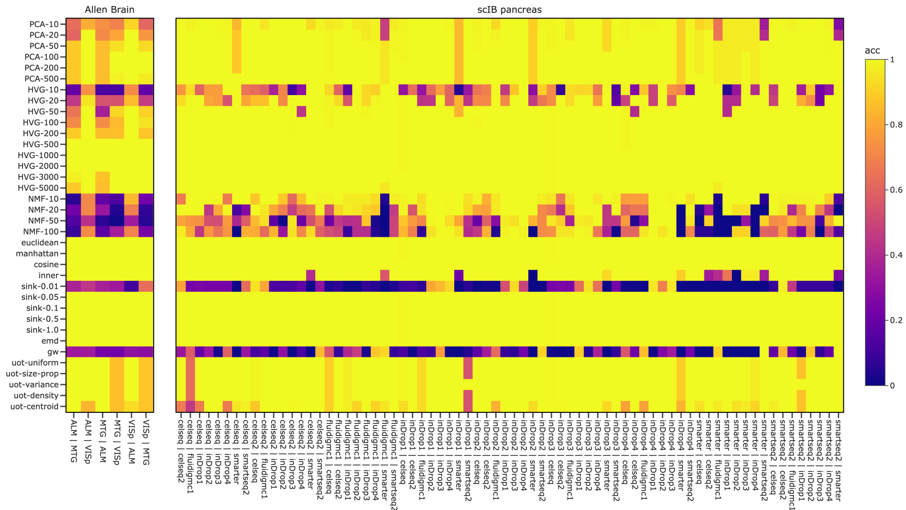


Supplementary Figure 2: Ablation study of RefCM hyperparameters across Allen Brain and scIB pancreas benchmark datasets. Heatmaps showing pairwise cell type annotation accuracy for different parameter configurations. Rows are grouped by three categories. (1) Embedding method and dimensionality: PCA, highly variable genes (HVG), and non-negative matrix factorization (NMF), each tested across varying feature dimensions; (2) Ground cost metric: Euclidean, Manhattan, cosine, and inner product distances for computing cell-to-cell costs; (3) Optimal transport solver: Sinkhorn with regularization parameter $\varepsilon \in \{0.01, 0.05, 0.1, 0.5, 1.0\}$, exact Earth Mover's Distance (EMD), Gromov-Wasserstein (GW), and unbalanced OT (UOT) with different mass weighting functions (uniform, size-proportional, variance-weighted, density-weighted, and centroid-weighted). Columns represent individual annotation tasks within each benchmark dataset. Note that Sinkhorn with $\varepsilon = 0.01$ exhibits poor performance due to numerical instability at low regularization values.

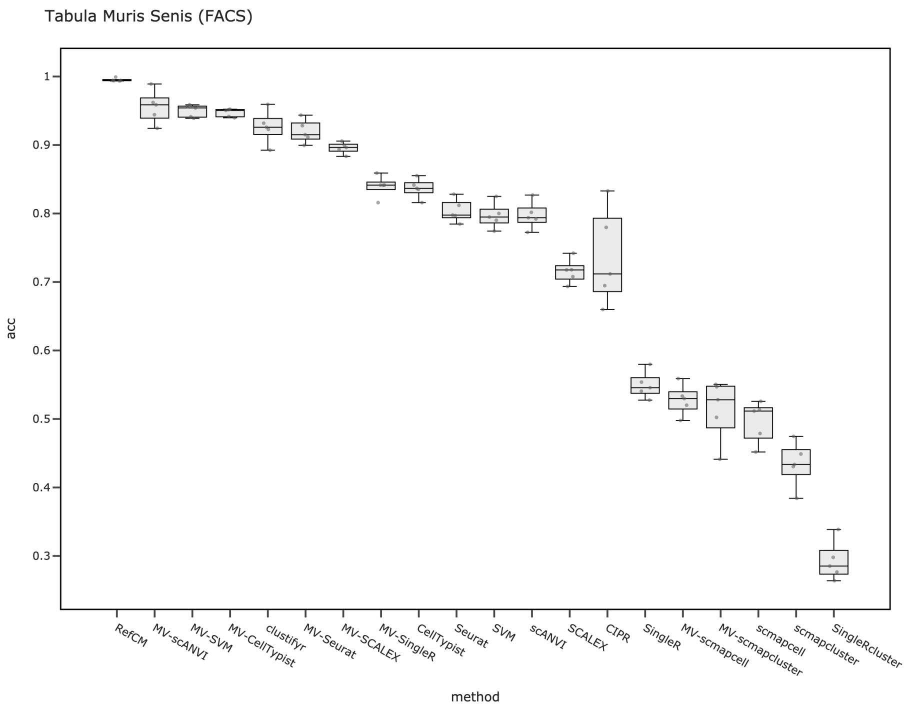


Supplementary Figure 3: Annotation accuracy across methods on Tabula Muris Senis (FACS). Each box summarizes accuracy over group-$k$-fold cross-validation splits (grouped by mouse ID). Methods ordered by descending median accuracy.
